# SAPTICoN, a robust no-code pipeline to analyze [plant] single cell transcriptomics data sets

**DOI:** 10.64898/2026.03.25.714156

**Authors:** Clément Pichot, Marion Verdenaud, Amandine Sandri, Gabrièle Adam, Etienne Delannoy, Pierre Hilson

## Abstract

Single-cell transcriptomic (SCT) analysis is essential for resolving cellular heterogeneity and uncovering the molecular foundations of development, physiology, and environmental responses. Despite its increasing importance, robust, reproducible, and broadly applicable SCT analytical tools remain largely restricted to well-annotated animal systems. This limitation poses significant challenges for biologists working on non-model species and poorly characterized tissues, where gene annotation is sparse and computational expertise is often limited.

We developed a stable, end-to-end SCT analysis pipeline designed to be accessible to biologists with little training in bioinformatics and applicable to species and tissues with limited genomic annotation. Built on the Seurat framework and complemented with additional tools, the pipeline supports any organism by automatically generating R-compatible annotation packages from basic genome files. It integrates standardized workflows for data preprocessing, quality control, clustering, biomarker identification, and gene set enrichment analysis within a fixed Snakemake framework, ensuring high reproducibility. By minimizing coding requirements, the pipeline enables rigorous, biologically informed SCT analyses across diverse experimental systems.

## INTRODUCTION

Single-cell transcriptomics (SCT) approaches are now well established to investigate molecular and cellular heterogeneity within sampled tissues or even entire organisms, based on RNA-seq libraries generated from isolated cells or nuclei (Zheng et al., 2017; Cao et al., 2017; Brown et al., 2024).

As in other phyla, multiple SCT studies have already described the role of individual cells in plant tissues (Yu et al., 2023). To cite a few, experiments capturing the transcriptome of a few thousand cells provide a useful overview of the role of specific cells and of mechanisms involved in tissue organization and differentiation, for example in root tips of Arabidopsis (Zhang et al., 2019; Denyer et al., 2019; Jean-Baptiste et al., 2019; Ryu et al., 2019; Shulse et al., 2019; Wendrich et al., 2020; Kim et al., 2021; Shahan et al., 2022; Han et al., 2024; Liu et al., 2024) and other plant species including crops (Zhang et al., 2021; Guillotin et al., 2023).

In parallel, a wide range of software packages have been developed in recent years to analyze large and complex SCT data sets, containing the transcriptome signatures of up to millions of cells. Most tools focus on a given step in the analytical workflow, for example data pre-processing (Zheng et al., 2017), cell clustering (Zhang et al., 2023), *de novo* cell marker detection (Hu et al., 2023 Hao et al., 2024), annotation cell population annotation (Aran et al., 2019; Clarke et al., 2021; Yang et al., 2022; Ji et al., 2023; Dong et al., 2024; Hou and Ji, 2024), developmental trajectory analysis (Cao et al., 2019) or topic modeling and annotation (Carbonetto et al., 2021; 2023). Some packages aggregate multiple functions, from the initial preprocessing and quality control steps, to later analyses and result displays, as singleCellExperiment (Amezquita et al., 2019) and Seurat (Hao et al., 2024).

But the choice between these different tools and their implementation is not trivial, especially for biologists without bioinformatics or coding expertise. Software and programming languages are regularly updated, and the transition between versions is far from seamless for a non-specialist. Even when their consecutive steps are linked, most computational workflows are not automated and slight configuration differences can alter the results.

Furthermore, existing SCT analytical pipelines are often built for species with well annotated genomes supported by a large community of users (Garcia-Jimeno et al., 2022; Cao et al., 2023) and it may be technically challenging to adapt them to species for which only custom genome annotations and gene function lists are available.

Another common hurdle for SCT neophytes is that there is no default method for the optimization of the clustering of cell populations based on their transcriptome. Yet, this step is essential to obtain results that faithfully represent the complexity of the characterized tissues and the relative representation of the distinguishable cell types, or that are corrected for potential bias inherent to the analysis (e.g. applied technology or experimental design). Finally, most of the available tools and pipelines have been developed and optimized for animal models, that may not be applicable to data characterizing cells from plants or other remote species (Panstruga et al., 2023).

It is therefore relevant to provide biologists with an up-to-date, automated and user-friendly tool to prepare raw SCT data for downstream analysis and to assist them answer their favorite biological questions. Such a tool should be designed as a robust analytical framework that integrates all required functions.

Here we describe a pipeline for SCT data analysis, named SAPTICoN, compatible with any annotated genome and without *a priori* about the structure and complexity of the input data (single cell (sc)RNA-seq or count tables). SAPTICoN is built as an automated and parallelized workflow that integrates the reference Seurat software with additional tools. It includes quality control of raw and pre-processed single cell (sc)RNA-seq data, downstream cell clustering, the study of *de novo* cell markers, and the analysis of expression profiles for genes of interest. In particular, SAPTICoN offers a module specifically designed to optimize cell clustering parameters that combines two unsupervised approaches. Users can thus rapidly compare different clustering configurations, test a range of thresholds for key parameters, and investigate whether downstream results reflect prior knowledge or expectations about the biological system under study. For example, are clusters adequately representing cell populations expected in the samples and are some known genes properly marking specific cell types? The SAPTICoN seamless package provides biologists with a reliable and comprehensive *in silico* knowledge basis to further pursue the study of their biological system. In fact, launching the pipeline involves filling out two files: the configuration file and the launch file, which greatly simplifies the process of launching the pipeline.

## RESULTS

### Pipeline structure

#### Framework

Our goal was to build a user-friendly tool that guarantees reproducible processes (Figure 1). SAPTICoN is launched via a single launching file that implements (i) Conda (Anaconda, Inc., 2017) to install the necessary resources and the work environment and (ii) Snakemake (Mölder et al., 2021) to organize the various analyses into configurable rules. Snakemake determines the dependencies between successive steps so that each of them is executed in the right order and only when necessary. Within the limit of available memory and processors, Snakemake also manages the execution of steps in parallel and the pipeline is thus adapted with either local (PC, laptop) or remote (server clusters) environments. This framework is flexible and new features can easily be integrated to add new functions.

**Figure 1.**
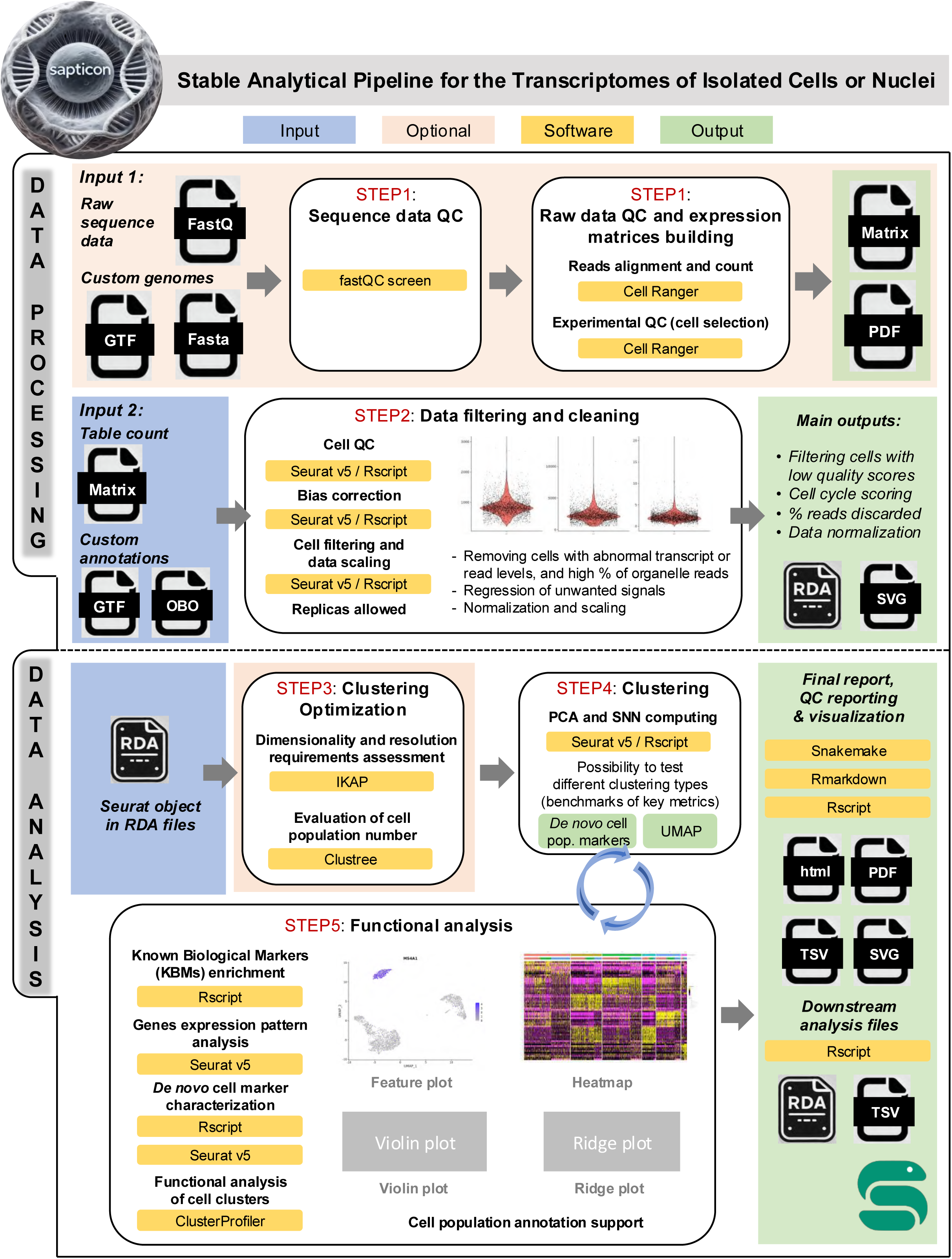
SAPTICoN workflow overview.

SAPTICoN is based on the following software for SCT data analysis, all working in R: Seurat v5 (Butler et al., 2018; Hao et al., 2024), arguably the most popular package including multiple functions; IKAP, that explores SCT data to find which numbers of cell clusters best represent populations characterized by differentially expressed genes (DEGs) (Chen et al., 2019); Clustree, an visualization tools to track the relationship between cell clusters according to varying resolution (Zappia and Oshlack, 2018); and clusterProfiler, designed for gene functional annotation and enrichment result analysis (Wu et al., 2021).

#### Input

Different files can be provided to start the analysis: raw fastq sequencing data or pre-computed expression matrix. They may correspond to a single or multiple samples, including replicates or not.

#### Output

At the end the pipeline provides a summary and an explanatory report for all the analyses performed. Each run produces plots and tables necessary for the biologist to interpret the data and, if relevant, to undertake additional analyses with the resulting Seurat object. This object records all the single-cell count data and metadata that are produced at each stage of the process. The SAPTICoN pipeline consists of several major parts, subdivided into different Snakemake rules. The next sections present an overview of the methods implemented in the logical order when starting a new analysis (Figure 1), but they do not explain in depth the rationale of each algorithm or statistical approach that are to be found in the original references.

### Data processing

#### From raw sequencing data to expression matrices (Input 1 / STEP1)

Most SCT technologies (e.g. Chromium, 10x Genomics; Quantum Single Cell RNA, ScaleBiosciences) produce large raw RNA-seq data files that are processed with the fastQC tool (Wingett and Andrews, 2018) to control the quality of the sequencing reads. CellRanger v7.1 software (Zheng et al., 2017) maps these to reference genomes, those of the nucleus and organelles in the species under study. A first count table (gene reads per cell) is then analyzed with another CellRanger module to discard putative cells with very few associated reads that are likely artefacts. A QC report is produced for the experiment, together with the expression matrix for each sample.

#### Data filtering and cleaning (Input 2 / STEP2)

Next, the pipeline further refines the selection of cells to be included in the analysis according to various criteria and thresholds (Seurat v5). Outlier cells with abnormal (high or low) numbers of transcripts or reads may be artefacts. A high ratio of organelle reads indicates a default of cell integrity.

Some samples may also require the removal of inherent biases by statistical linear regression, for example gene expression signatures linked to stress caused by plant cell wall digestion for protoplast isolation or to cell cycling in proliferating tissues.

This multi-step filtering process is specific to each data set. It ends with the removal of the discarded cells and of potential noise, yielding cleaned-up expression matrices. The filters and corrections applied are specific to each data set and depend on the experimental design. Finally, the count tables are normalized and scaled, and the data undergoes principal component analysis (PCA).

### Data analysis

In the SAPTICoN pipeline, the data analysis *per se* consists in three successive stages, hereafter listed as steps 3 to 5 (Figure 1). First (STEP3), the biologist searches the best compromise when choosing key parameters for the clustering of similar groups of cells. Such parameters are sometimes fixed by default or set with simple general rules. Our experience is that each data set is unique and deserves careful consideration to choose an appropriate clustering configuration because this initial step determines to a large extent the conclusions that will be drawn from the study. More information about selection of clustering parameters will be given in the next section. Second (STEP4), the pipeline builds the cell clusters based on the selected clustering parameters and provides different representations of the data, combining information relative to cell groups and gene expression patterns, as well as lists of markers specifying cells of interest. Finally (STEP5), the pipeline investigates whether groups of genes, whose expression pattern varies according to cell populations, provide valuable information about their role and the biological functions of the different cells.

#### Optimization of clustering parameters ***(STEP3)***

Prior to the clustering completed in STEP4, a value must be set for two parameters: (1) the number of principle components (nPCs) to include in the data dimensionality reduction; (2) the resolution (*r*) that sets the granularity of cell clustering (higher *r* values yielding more *k* clusters). Indeed, it is not possible to accurately predict the number of cell groups present in a biological system or to assign a priori a coherent cell annotation faithfully reporting its content. The proper values for clustering parameters also depend on experimental design (number of cells, sample treatments, technology to produce the SCT data, etc.). Four methods are included in the pipeline for parameter optimization.

**1. Elbow plot** (in Seurat). This graph plots the variation of the data explained by the successive PCs. The number of PCs to include are indicated by an inflexion point beyond which only minimal gain is obtained by additional PCs. The method is often judged imprecise and may not point to a clear “elbow” depending on the structure of the data (Figure 2A).
**2. JackStraw test** (in Seurat). This unsupervised method calculates the statistical significance of associations between the observed data and their systematic patterns of variation. This information can be used to estimate the number of PCs beyond which the association becomes non-significant (Figure 2B) (Chung and Storey, 2015).
**3. IKAP**. This other unsupervised method systematically browses through the number of top principal components (nPC) and the resolution (*r*) for Shared Nearest Neighbor (SNN) clustering (in Seurat) and identifies the combination(s) most likely to produce distinctive marker genes (Chen et al., 2019). IKAP first determines a range of PC numbers (PCmin, PCmax) and a maximum number of clusters (*k*max) to explore the parameter space. The algorithm then scans, for each given nPC, how the gap between observed and random data evolves as *r* increases (from fine to coarse grouping) assuming that a large gap indicates a shift towards a clustering that separates well major groups. Finally, based on DEGs identified for the top candidate combinations, decision trees are constructed to assess cluster differentiation, and the classification error is calculated as the weighted sum of cluster-level errors across multiple splits of the decision tree. The best combination has the lowest average classification error. IKAP provides a summary heatmap of the gap scores including marks pointing the best recommended combination and two good alternatives (Figure 2C).
**4. Clustree**. This tool enables the rapid visualization of the relationship between groups of cells across varying resolutions (*r*), for a given number of PCs (Zappia and Oshlack, 2018). The analysis is presented as a tree in which the edges show overlaps between clusters (nodes). Appropriate resolutions correspond to the largest number of *k* clusters (highest *r*) for which the relationships with lower-order clusters (lower *r*) are local and distinct, rather than unstable. In the tree, the clusters’ relationships can be viewed associated with a metric that evaluates mathematically cluster stability across increasing resolution (SC3 method; Kiselev et al., 2017) (Figure 2D) or simply browsed (Zappia and Oshlack, 2018) (Figure S1).

**Figure 2.**
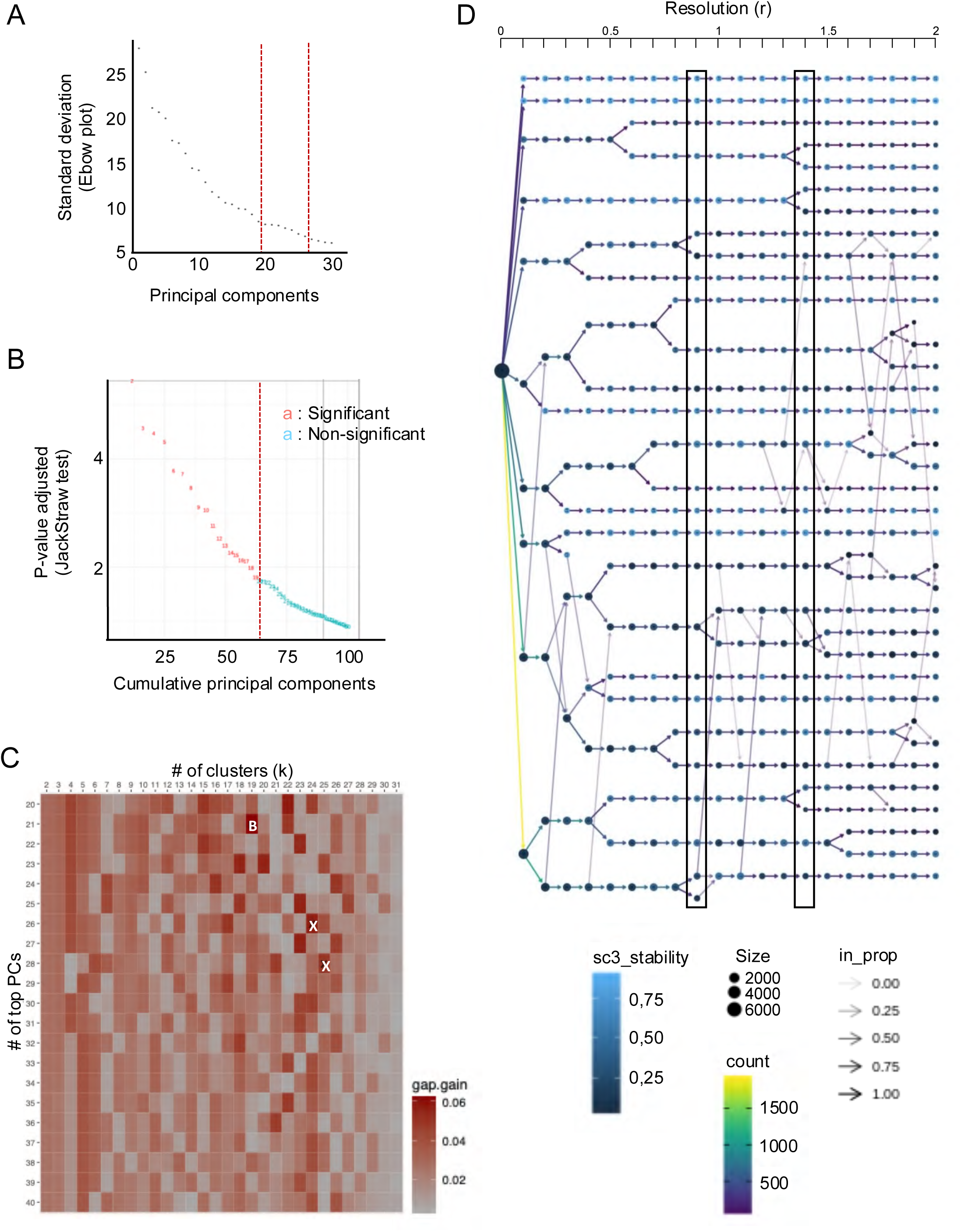
SAPTICoN clustering optimization. All panels illustrate the analysis of the single-cell transcriptomics reference data published by Sharan et al. (2022). (A) Elbow plot, generated in Seurat. Curve inflection indicates that principal components (PCs) from 20th on explain a marginal portion of data variation. (B) JackStraw test plot, generated in Seurat. Red numbers represent significant PCs that explain data variability, the last one being the 19th. (C) Heatmap representing the best clustering configuration as an output of the IKAP software analysis. B: best; X: alternatives. Red color depth represents gain gap. (D) Clustering tree generated with the Clustree software. Dot size represents the number of cells within each cluster. Dot color represents cluster stability as resolution (*r*) increases from 0 to 2, in increments of 0.1. Arrow color represents the number of cells common between affiliated clusters. Black box indicates the best clustering parameters.

While the Elbow plot or JackStraw methods provide an estimate of nPC that is often used for SCT data dimensionality reduction, we prefer to choose nPC, resolution (*r*) and cluster number (*k*) parameters as a compromise in line with both the IKAP and Clustree results, after verifying that slight variations around the chosen parameters do not affect cell clustering. Nevertheless, within SAPTICoN, each user has the choice to compare and use any of these methods to choose the clustering parameters that best fit the data. Furthermore, the pipeline offers the possibility to include prior knowledge in this optimization step (see Discussion).

#### Clustering (STEP4)

With Seurat, cells are clustered based on their transcriptome with modularity optimization techniques such as the Louvain algorithm or SML (Blondel et al., 2008) to iteratively group similar cells together. To visualize the resulting clusters, the cells are then projected into a low-dimensional space with dedicated dimensionality reduction approaches, such as uniform manifold approximation and projection (UMAP) (McInnes et al., 2018) or t-distributed stochastic neighbors’ integration (t-SNE) (van der Maaten and Hinton, 2008) (Figure 1).

#### Functional analysis (STEP5)

Once cell groups have been defined, the respective role of the different cells can be investigated based on their specific gene expression signatures. For this purpose, *de novo* cellular markers specific to one (or more) cluster are selected through differential analysis comparing the median expression of genes in that cluster to that of the other clusters (in Seurat, Hao et al., 2024) (Figure 1). Furthermore, SAPTICoN enables the user to match defined groups of cells with prior knowledge by performing a Fisher enrichment test that compares the list of corresponding *de novo* cell marker genes with lists of biological markers known to be specific of identified cell types (Known Biological Markers, KBMs), thereby pinpointing the likely cell type(s) grouped in certain cluster(s).

In addition, the expression profile of any marker gene (identified *de novo*, based on previous knowledge, etc.) can be investigated across cell clusters in different ways: feature plots, to assess gene expression per cell as colors on the projected cell map; violin or ridge plots, showing expression level distribution for a given gene per cell, across the clusters; heatmaps, for gene expression as levels in cells arranged per cluster.

Finally, with the aim to assign functions and roles to particular groups of cells, gene set enrichment analyses (GSEA) are performed based on lists of *de novo* marker genes that characterize these groups, leading to a refined cell annotation that blends the SCT results with prior data about the tissues and biological system under study (Figure 1).

### Fixing clustering configuration and cell annotation

As previously mentioned, the parameters that enable the extraction of original quality information from SCT data sets are not trivial to establish. Their choice requires the confrontation of the pipeline’s preliminary results with prior biological knowledge. The clustering optimization embedded in the pipeline (STEP3) suggests a small number of potentially appropriate configurations (nPC, *r* resolution*, k* number of clusters) based on the intrinsic structure of the data to further analyze through clustering (STEP4) and functional analysis (STEP5). The main task for the biologist is thus to examine which of these configurations best reflects what is already known by comparing results of a few configurations. This iterative process provides answers to the following questions. Do specific groups of cells express expected known biological markers (KBMs)? Do key biological processes known to take place in such cell populations match with their associated *de novo* markers? Are all related cells grouped in one cluster or in closely mapped clusters? Do the clusters appropriately represent the cell type diversity expected in the sample?

The configuration that most satisfactorily answers these questions is then fixed and the cells clusters are annotated accordingly. At the end of the SAPTICoN automated pipeline, a Seurat object is built with the corresponding metadata and a complete, explanatory and detailed report of the various results is provided in markdown format (Figure 1).

This Seurat object can be further exploited for more advanced downstream analyses such as trajectory inference (Cao et al., 2019), TOPIC modeling (Dey et al., 2017), RNA velocity (La Manno et al., 2018) or additional differential expression analyses for clusters of particular interest to the user’s biological questions, for example in the scPlant framework (Cao et al., 2023). In our view, such analyses are complex and difficult to automate for technical reasons and because they depend on data heterogeneity. We have chosen not to implement them in the pipeline to maintain its ease of use.

### Benchmark

To test the SAPTICoN pipeline, we reanalyzed SCT data generated by Shahan et al. (2022) to study the cell populations making up the tip of young wild-type primary roots in *Arabidopsis thaliana*. This organ has a simple radial symmetry maintained through indeterminate growth and its development is well described with all main cell types and division events leading to root tissue differentiation precisely mapped. In brief, Shahan et al. aggregated their own original data with those of several independent reports and build a comprehensive Arabidopsis root cell atlas. For this purpose, cell labels were established by combining different sources of information: (i) developmental stage assignments based on curated marker genes; (ii) expression profiles extracted from published spatial transcriptomic resources; (iii) marker-based annotation using known cell-type genes; (iv) correlation-based annotation to previously profiled transcriptomes; and (v) Index of Cell Identity (ICI) values that measure similarity to predefined signatures.

Our goal was to compare the Reference clusters obtained by Shahan et al. with those produced with minimal intervention by the SAPTICoN pipeline. For clarity, the SAPTICoN-driven analysis is detailed here below according to the main steps of our workflow (Figure 1).

***Data processing*** (STEP1 and STEP2). The CellRanger pipeline’s quality control retained a total of 6,779 cells with an average of 3,576 genes per cell (min=820; max=10,982) and an average of 20,381 UMIs per cell (min=1,359; max=20,381) (Figure S2). These cells are not exactly identical to the 6,433 cells selected by Shahan et al. from the same initial count table for further analysis, but they are largely overlapping with 6,433 common cells. The number of reads associated with organelle genomes was very low (mitochondrion <5%; chloroplast <0.05%) and was considered negligible (Figure S2) as such ratios indicate a cell population of sufficient quality for downstream analysis.

***Optimization of clustering parameters*** (STEP3). IKAP selected clustering parameters of 21 PCs with a resolution (*r*) of 1.9 for its best configuration, its two alternatives being 26 PCs/*r*=2.4 and 28 PCs/*r*=2.5 (Figure 2C). Clustree selection is 25 PCs and *r* between 0.9 and 1.4 (Figure 2D; Figure S1). The corresponding tree shows that cluster filiation is relatively stable starting at a resolution of 0.9 up until 1.8, with a SC3 stability, a score that measures how well a cluster remains consistent and retains largely the same cells when the clustering resolution is changed, greater than 0.5 for most branches in that range (Figure 2D). IKAP- and Clustree-selected nPC values are similar to those indicated by the simpler optimization methods: the pipeline-produced Elbow plot shows a marked plateau between 20 and 30 PCs (Fig. 2A); the JackStraw graph determines that 60% of the data variability is significantly explained by the first 19 PCs (Fig. 2B).

***Clustering*** (STEP4). At this stage, the pipeline data processing retained 6,433 cells, the associated expression matrix has been scaled and normalized and the data was subjected to PCA. We had now to settle on a final clustering configuration in the range of those suggested in the previous SAPTICoN step: between 19 and 28 PCs and a resolution around 1.0. To that end, we compared clustering results with nPC=19 or 25, and *r*=0.5 or 1.0. As expected, different resolutions slightly affected the number of clusters, but the structure of the UMAP projections were almost identical (Figure S3).

We chose the clustering parameters nPCs=25 and *r*=1.0 as a good compromise, yielding 26 Seurat clusters (Figure 3A). In Shahan et al. (2022), cells in the Reference data were grouped in 64 Seurat clusters (Figure 3B). The clustering based on the SAPTICoN “compromise” configuration thus contains 2.5 times fewer clusters (Figure 3C), with sufficient cell segregation to represent the presumed cell populations, suggesting that a simpler cell distribution may better reflect the population diversity in the sampled root tips and that the repartition of the cells in the 64 Reference clusters may be overfitted.

**Figure 3.**
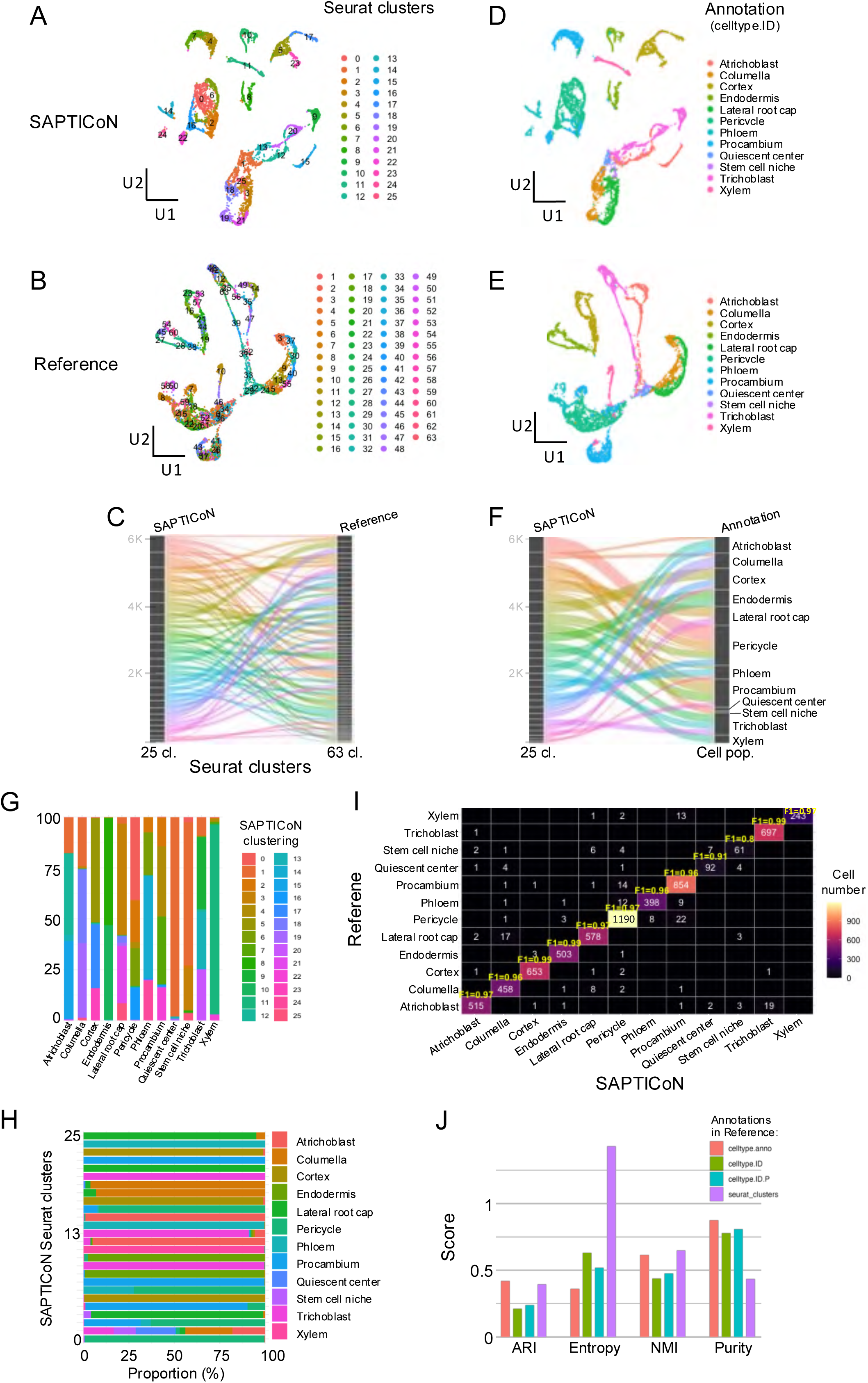
SAPTICoN versus Reference analysis of published Arabidopsis root cell data. (A) UMAP visualization of the cell populations characterized by Shahan et al., 2022, after data pre-processing and Seurat clustering with the SAPTICoN pipeline. (B) UMAP visualization of the Reference data set as pre-processed and clustered with Seurat by Shahan et al. (2022). (C) Cell-based correspondence between clusters defined in the SAPTICoN and Reference data sets. (D) Cell representation in the SAPTICoN UMAP, after transfer of the celltype.ID annotation from Shahan et al. with Seurat’s anchor-based label transfer functions. (E) Cell representation according to the original annotation in the Reference data set. (F) Cell-based correspondence between SAPTICoN-defined clusters annotations. (G) Cumulative bar plot of SAPTICoN-defined Seurat cluster cell proportions (%) per annotation. (H) Cumulative bar plot of annotated cell proportions (%) per SAPTICoN-defined Seurat cluster. (I) Confusion heatmap of comparing cell annotations between the SAPTICoN and Reference data sets. F1 scores are calculated based on annotations of cells common to both sets to assess the concordance between the two pipelines. (J) Comparison of clusters and annotations produced by SAPTICoN, based on concordance metrics (ARI, entropy, NMI, Purity), for four Reference annotations. Orange: celltype.anno; green: celltype.ID; turquoise: celltype.ID.P; purple: initial Reference Seurat-defined clusters. The Purity metric reflects the dominant proportion of a single cell type per cluster: a high value indicates well-separated and homogeneous clusters.

***Pipeline validation*** (STEP5). To further investigate this issue, we transferred multiple cell-type annotation schemes from the Reference data set of Shahan et al. to the SAPTICoN data set using Seurat’s anchor-based label transfer functions. This procedure identifies anchors between the Reference and query SAPTICoN data sets and transfers labels based on cell transcriptome similarity, returning per-cell prediction scores that were used for stringent confidence filtering to reduce mislabeling of out-of-reference states.

To ensure that our conclusions were not based on a single labeling system, we evaluated the concordance between labels transferred to cells in the SAPTICoN clusters and four views provided by Shahan et al. (2022): (1) cellType.anno, fine annotation labels with based on developmental signatures, (2) cellType.ID, a coarser identifier (illustrated in Figure 3D to F) based on other cell type marker genes, (3) cellType.P.ID, based on probability-weighted labels derived from Seurat cell prediction scores; and (4) the cell clusters as identified in the initial 64 Reference Seurat clusters (Figure 3B).

The cellType.anno and cellType.ID transferred cell labels associated to the SAPTICoN Seurat clusters were consistent with expected cell types and adequately characterized the different defined clusters, with proper boundaries reflecting the Reference taxonomy. These observations, taking into consideration multiple label granularities and integrating of cell-based confidence scores that minimize taxonomy- or threshold-dependent artefacts, confirm the relevance of the “compromise” clustering from a biological point of view (illustrated for transferred cellType.ID labels in Figure 3G and H).

### Reference and SAPTICoN cell annotations are essentially overlapping

To quantitatively assess the reciprocity of the Reference and SAPTICoN clustering data sets, we analyzed the number of cells shared between them with several metrics (see Methods). The annotations of the SAPTICoN clusters have F1 scores >0.97 for 10 of its 12 cell populations (Figure 3I). The *Adjusted Rand Index* (ARI) which shows how cells are classified according to annotation shows average reciprocity (Figure 3J). As expected, the Reference Seurat clusters have the highest entropy score, confirming greater diversity of annotations per cluster before re-annotation of the cell populations, and the over-clustering of the reference data set. The celltype.anno and celltype.ID.P annotations show good performance in NMI and Purity, indicating a better overall match with the reference labels for the three types of annotation tested (Figure 3J).

Finally, we analyzed the impact of the transformations performed in the SAPTICoN pipeline on gene expression, in general, or with a focus on the most variable genes (HVGs), the *de novo* markers and cell type markers. Expression profiles of genes marking specific root cell types or tissues are equally illustrative in both UMAP projections (Figure 4A; Reference and SAPTICoN in upper and lower panels, respectively) and feature plots (Figure 4B; compare to Figure 1E in Shahan et al., 2022). The distribution of gene expression variance relative to the reference is preserved in the SAPTICoN data set, indicating that the pipeline does not introduce any obvious bias (Figure 4C). This observation agrees with the overlap of the reference and SAPTICoN cell population projections according to the two top components, following the PCA of either data set based on the 3,000 most variant genes (HVGs) (Figure 4D). Also, *de novo* cell markers associated to clusters defined in either data set are very similar (Fig. 4E).

**Figure 4.**
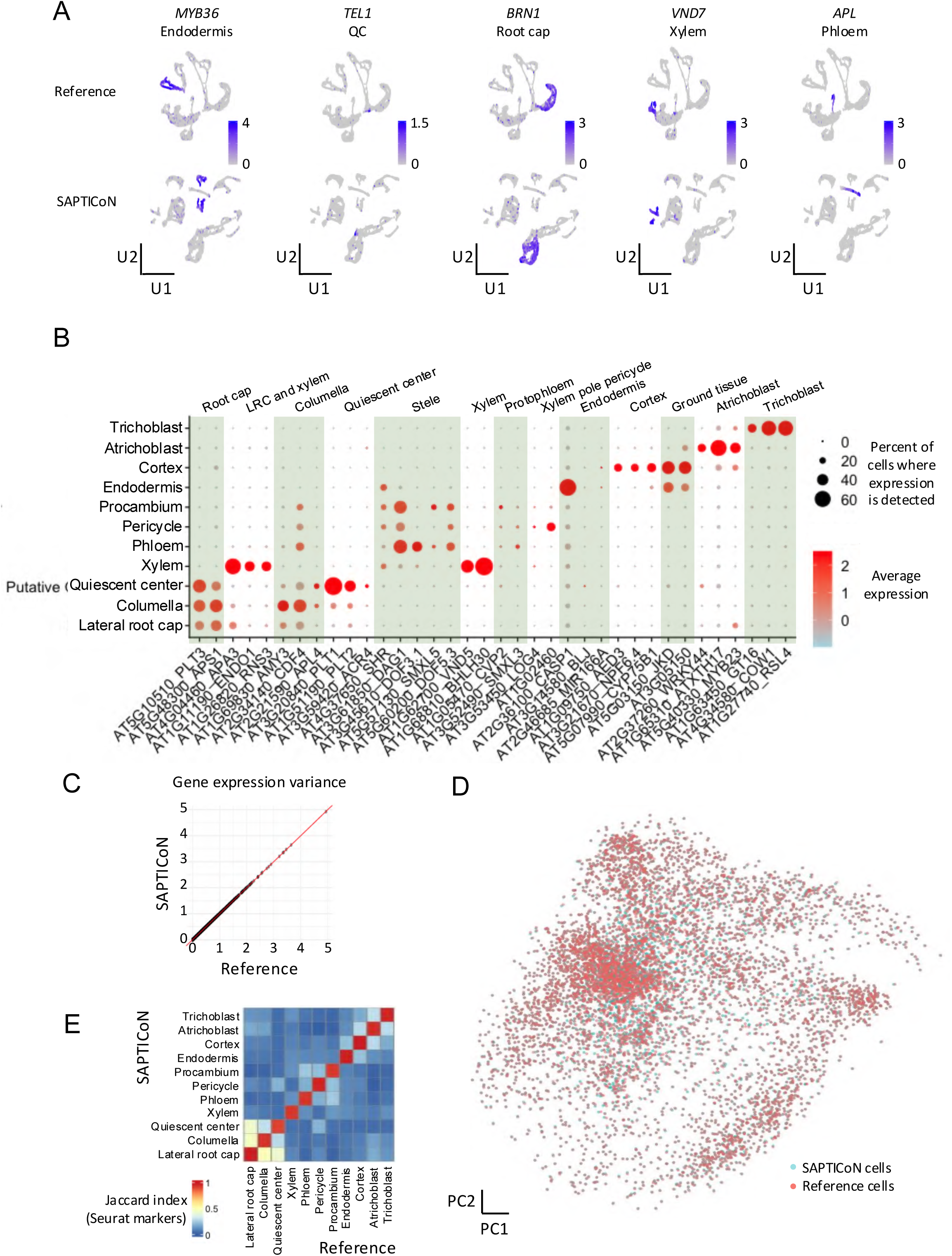
Gene expression marker profiling. (A) Expression of known biological markers (KBMs). Blue scales represent normalized expression levels. *MYB36*, AT5G57620; *TEL1*, AT3G26120; *BERN1*, AT1G33280; *VND7*, A1G71930; *APL*, AT1G79430. (B) Average expression of KBMs per cell population. Ratio of cells is proportional to dot size and expression shown according to color scale (compare to Figure 1E in Shahan et al., 2022). (C) Gene expression variance comparing the SAPTICoN and Reference data. (D) PCA cell projection according to gene expression in SAPTICoN and Reference data sets. (E) Jaccard index of the Seurat *de novo* markers per cell population in the SAPTICoN and Reference data sets.

In conclusion, our results indicate that the methods implemented in the SAPTICoN pipeline does not distort expression profiles nor does it introduce analytical bias. Our benchmark study shows that the pipeline captures significant biological structures that mostly overlap with those defined in the Reference analysis. Furthermore, the initial SAPTICoN Seurat clustering contained fewer clusters fixed before cell annotation, and thereby simplified the completion of this crucial step.

## DISCUSSION

### The drive to develop SAPTICoN

The pipeline was designed for biologists who wish to process their original SCT data sets and have a theoretical understanding of the required analytical methods, but lack the expertise to manage consistently the nitty-gritty details of their implementation in the ever-evolving R environment. For them, the learning curve is steep and scripts that worked one day may be disabled the next or yield inconsistent results due to changing algorithm versions.

The locked and validated SAPTICoN pipeline ensures reproducibility while enabling the experimenter to pick key parameters that are best adapted to their samples. The package assembles tried-and-tested methods combined for their respective advantages.

### Matching clustering options with prior knowledge

Here, we illustrated the usefulness of the integrated pipeline through the analysis of a data set initially produced and published by Shahan and coworkers (2022) that characterizes the different cell populations involved in the development of the Arabidopsis primary root.

We focused in particular on the IKAP and Clustree unsupervised methods designed to choose pertinent values for clustering parameters (number of principal components included [nPC], resolution [*r*], number of clusters [*k*]). Based on the tested data set, we showed that the IKAP and Clustree nPC and *r* ranges yield cell clusters that correspond well with the cell labels resulting from the careful annotation of Shahan et al. With no other information than the SCT data itself, the main root cell types were properly identified, avoiding overfitting that may produce supernumerary clusters and complicate the initial analysis of the results.

In addition, SAPTICoN facilitates in-depth analysis of all considered clustering configurations. Because it is automated, the SAPTICoN pipeline can (i) systematically analyze and display the expression profiles of markers associated to each generated cell cluster, and (ii) functionally characterize these clusters through GSEA, this being performed for all clustering configurations of potential interest. Thereby, when key cellular markers are already known, it is straightforward to complete clustering optimization by checking whether such KBMs are in fact listed as *de novo* cellular markers for specific cluster(s) and whether markers of different cell populations are significantly enriched in distinct clusters. This level of detail allows users to directly compare prior knowledge with the subtle effects that variations in clustering parameters may have on cell annotation labels.

### Comparison with other tools

Other integrative SCT analytical pipelines are available to biologists that share functions between themselves and with SAPTICoN (Guo et al., 2015; Zheng et al., 2017; Parekh et al., 2018; Hoek et al., 2021; Moreno et al., 2021). However, most focus on specific tasks and the few automated pipelines do not include both preprocessing functions and tools for comprehensive biomarker identification, gene function analyses and cell annotation (Table 1 and weblinks therein). A software similar to SAPTICoN is scPlant, a sound framework for the complete analysis of single-cell data initially designed for plants, but it is not automated and thus requires a certain command of programming languages and bioinformatics (Cao et al., 2023) (Table 1). Another tool similar in structure is the Bollito software package (García-Jimeno et al., 2022). Bollito enables automation and parallel analyses, includes the various common steps through to the functional analysis of specified cell populations, and it is based on the Snakemake workflow management system and a Bioconda package repository (Table 1). In comparison, SAPTICoN adds methods to fine-tune the clustering parameters.

**Table 1.** Overview of pipelines for SCT data analysis.

|  | SAPTiCoN | scPlant | Bollito | WASP | SCiAP | iCellR | zUMI | SINCERA |
| --- | --- | --- | --- | --- | --- | --- | --- | --- |
| **Pipeline core** |  |  |  |  |  |  |  |  |
| Automation | ✓ | ✗ | ✓ | ✓ | ✗ | ✗ | ✗ | ✗ |
| Parallelized | ✓ | ✗ | ✓ | ✓ | ✗ | ✗ | ✗ | ✗ |
| Dependency management | ✓ | ✓ | ✓ | ✓ | ✗ | ✗ | ✓ | ✗ |
| **Data pre-processing** |  |  |  |  |  |  |  |  |
| RNA-seq raw data QC | ✓ | ✗ | ✓ | ✓ | ✗ | ✗ | ✓ | ✗ |
| Reads alignment & count | ✓ | ✗ | ✓ | ✓ | ✓ | ✓ | ✓ | ✗ |
| Experimental QC | ✓ | ✗ | ✗ | ✓ | ✓ | ✗ | ✓ | ✓ |
| Retained data QC | ✓ | ✓ | ✓ | ✓ | ✗ | ✗ | ✓ | ✓ |
| Cleaning & normalization | ✓ | ✓ | ✓ | ✓ | ✓ | ✓ | ✗ | ✓ |
| **Marker gene analyses** |  |  |  |  |  |  |  |  |
| Clustering optimization | ✓ | ✗ | ✓ | ✗ | ✗ | ✗ | ✗ | ✗ |
| Clustering | ✓ | ✓ | ✓ | ✓ | ✓ | ✓ | ✗ | ✓ |
| *De novo* marker identity | ✓ | ✓ | ✓ | ✓ | ✓ | ✓ | ✗ | ✓ |
| KBM enrichment analysis | ✓ | ✗ | ✗ | ✗ | ✗ | ✗ | ✗ | ✗ |
| Marker gene profile assess. | ✓ | ✗ | ✗ | ✗ | ✗ | ✗ | ✗ | ✗ |
| **Functional analysis and cell annotation** |  |  |  |  |  |  |  |  |
| BSgenome packaging | ✓ | ✗ | ✗ | ✗ | ✗ | ✗ | ✗ | ✗ |
| GSEA analysis | ✓ | ✓ | ✓ | ✗ | ✗ | ✗ | ✗ | ✗ |
| Trajectory inference | ✗ | ✓ | ✓ | ✗ | ✓ | ✗ | ✗ | ✗ |
| TOPIC analysis | ✗ | ✓ | ✓ | ✗ | ✗ | ✗ | ✗ | ✗ |
| RNA velocity | ✗ | ✗ | ✓ | ✗ | ✗ | ✗ | ✗ | ✗ |
| **Model study application** | ----- Plant ----- |  | ----- Animal ----- |  |  |  |  |  |
| **Reference year** | 2024 | 2023 | 2021 | 2021 | 2021 | 2020 | 2018 | 2015 |
| **URL** | (1) | (2) | (3) | (4) | (5) | (6) | (7) | (8) |

(1) https://forge.inrae.fr/bcr/; (2) https://github.com/compbioNJU/scPlant; (3) https://github.com/cnio-bu/bollito; (4) https://github.com/andreashoek/wasp; (5) https://github.com/ebi-gene-expression-group/container-galaxy-sc-tertiary; (6) https://github.com/rezakj/iCellR; (7) https://github.com/sdparekh/zUMIs; (8) https://research.cchmc.org/pbge/sincera.html

### Automated formatting of gene annotations as R-compatible packages

Another common hurdle is that SCT analytical software generally assume that the genome data for a species of interest are readily available or fully referenced in dedicated databases. But that may not be the case and some packages may therefore not be fit for purpose because high-quality gene structural and functional annotations are required for accurate enrichment analyses (e.g. through GSEA) that point out biologically meaningful pathways and processes associated with particular cell types.

In the R environment, gene annotations are generally formatted and accessed as BSgenome objects (Pagès, 2024), as in the clusterProfiler software (Wu et al., 2021) integrated in the SAPTICoN pipeline, to perform GO (Gene Ontology) and KEGG (Kyoto Encyclopedia of Genes and Genomes) term enrichment analyses.

However most sequenced and annotated plant genomes are not available as pre-built BSgenome objects (for example in the UCSC Genome Browser; https://genome.ucsc.edu). Furthermore, the construction of such custom packages requires advanced knowledge of R/Bioconductor, significant time and in many cases the involvement of repository administrators. This process can thus significantly delay analyses even for experienced bioinformaticians.

To solve these hurdles, SAPTICoN implements an automated rule that builds a fully functional custom BSgenome package directly from the reference genome and protein sequence FASTA files provided to the pipeline. This automated step speeds up downstream analyses and facilitates access to high-quality functional annotations, enabling researchers to perform comprehensive GSEA on non-model species with minimal manual intervention. In fact, sharing pre-packaged BSgenome objects could be a useful standard across SCT communities dedicated to a common species, and SAPTICoN provides R tools for such practice.

To conclude, we hope that the SAPTICoN pipeline will be helpful to a wide-range of biologists who wish to study of their biological system through single-cell transcriptomics approaches, but do not have advanced bioinformatics skills. It is freely available at GitHub (https://forge.inrae.fr/bcr/SAPTiCon) for non-commercial use. The software includes comprehensive documentation that guides users through pipeline installation and explains each analytical step in practical, accessible terms.

## METHODS

### Data sets

For the single cell transcriptome-based data analysis, scRNA-seq data were retrieved from GEO data repository GSE152766 (Shahan et al., 2022).

### SAPTICoN implementation

The SAPTICoN pipeline is primarily implemented in R, leveraging its robust statistical and graphic capabilities. We opted for the Seurat object (Hao et al., 2024) to manage the underlying data structures of single-cell data. In addition, the pipeline integrates R packages specifically developed for single-cell transcriptome analysis such as IKAP (Chen et al., 2019), Clustree (Zappia and Oshlack, 2018). The pipeline organization is managed with Snakemake (Mölder et al., 2021) for efficient workflow execution and management. Automatic installation of environments for dependency requirements is handled through Conda (Anaconda, Inc., 2017), streamlining the setup process. This combination allows for seamless integration and reproducibility across different computational environments. Finally, prior installation of the Cell Ranger software (10X Genomics) for data preprocessing is required if this method is used.

### Preprocessing of Sequencing Data and quality control

Quality control of raw sequencing data was assessed with fastQC screen (Wingett and Andrews, 2018). As the data set was built using 10X Genomics technology, we have implemented CellRanger v7.1 (Zheng et al., 2017) to assess the experimental quality control and available cell selection, the mapping, and expression matrix building.

### Pre-processing and quality control of single-cell data

Gene expression count matrices are loaded into individual Seurat objects, or embedded into a single object in the case of technical replicates, using the Seurat R package v5.1.0 (Hao et al., 2024). Cells are selected for further analysis according to several criteria: expressed gene number and/or UMI count within specified ranges, percentage of mitochondrial gene reads below a given threshold, cell cycle scoring (according to G2/M and S phases) with Seurat functions. The data is then scaled for each sample using the *scaleData()* functions. Two methods are provided for normalization (Choudhary and Satija, 2022): *LogNormalize()* computes the relative expression of each gene (gene read count divided by cell read total), that is then multiplied by a scaling factor (10 000 by default) and log transformed. The *SCTransform()* models UMI counts using a regularized negative binomial regression, providing variance-stabilized and normalized data. Results are presented with different plots in the final report.

### Technical biases correction and cleaning

Following the initial quality control assessment, unwanted cells are discarded with user defined criteria: number of expressed genes and/or UMI counts and a maximum percentage of mitochondrial gene expression (superior to 5% by default). To minimize effects that may due to technical issues, for example contamination by organelle RNAs or stresses associated with enzymatic cell wall digestion, apply a regression step based on gene characteristic to effects such as genes that are linked to (i) cell cycle, (ii) mitochondrion/choloroplast-related processes, or (iii) cell protoplasting via the *scaleData()* function, setting the vars.to.regress = “geneList” parameter to the results from *cellCycleScoring()* or only from the list of genes of interest if bias is not associated with the cell cycle.

### Clustering optimization

Following PCA of the expression matrix, the evaluation of optimal parameters for clustering single-cell data is carried out with four different approaches. The Seurat *ElbowPlot()* function evaluates the number of PCs to include in dimension reduction as this heuristic method generates an ’Elbow plot’ that ranks principal components based on the percentage of variance explained by each of them in order to capture the majority of the true signal. The Seurat *JackStraw()* function evaluates the number of PCs to include as it determines the statistical significance of PCA scores (Chung and Storey, 2015). The IKAP software (Chen et al., 2019) investigates sets of cell clusters (clustering) by adjusting the number of top principal components (nPCs) and resolution (*r*) for Seurat SNN clustering. It selects a few candidates sets from all those examined, one of which is indicated as the best and most likely to produce distinctive marker genes using the default parameters. Finally, the Clustree tool (Zappia and Oshlack, 2018) evaluates the resolution (*r*) with the default parameters. It displays relationships as a tree linking clusters at different resolutions to assess whether cluster contents varies or not across *r* values. Calculated cluster similarity can also be shown in the tree (SC3 method; Kiselev et al., 2017).

### Cell clustering and dimensionality reduction

In Seurat, cells are clusters based on their PCA scores, either with the Louvain algorithm or SML (Blondel et al., 2008), with the optimized nPC and resolution values in the *FindNeighbors()* and *FindClusters()* functions, respectively. Uniform manifold approximation and projection (UMAP) (McInnes et al., 2018) or distributed stochastic neighbor embedding (t-SNE) (van der Maaten and Hinton, 2008) produce low-dimensional embeddings based on the user-definable upper principal components. The non-parametric Wilcoxon rank-sum test method, implemented with the Seurat function *FindAllMarkers()*, identifies the most highly variable genes (HVGs) across the entire cell population. By default, the number of HVGs included in further analyses is 3,000. This parameter can be defined by the user depending on the median number of genes detected within cells.

### Representation of gene expression signatures per cell groups

A function of Seurat visualization functions is used to represent gene expression profiles according to clusters (or other identity groups) by extracting data from the Seurat object: feature plots on map projection, *FeaturePlot()*; heatmaps, *DoHeatmap()*; violin plots, *VlnPlot()*; ride plots, *RidgePlot()*.

### Import of gene annotations

Gene annotations, such as GO and KEGG data, may be imported with the genome data (GTF and Fasta files). Alternatively, they can be supplied to SAPTICoN, with the annotation R package BSgenome (Pagès, 2024), to build and format the databases and R files required for downstream analysis. Alternatively, SAPTICoN can build in automatized manner the KEGG and GO databases with eggNOG v5 based on protein sequence fasta files (Huerta-Cepas et al., 2019).

### Gene enrichment analysis

Enrichment of GO terms linked to genes specific to groups of cells is measured with the gene set enrichment analysis (GSEA) function of the clusterProfiler package (Wu et al., 2021). Genes associated with significantly enriched terms are listed in table format.

### Evaluation of annotation reciprocity

Four cell-type annotation schemes were transferred from the Reference data set of Shahan et al. to our integrated data set using Seurat’s anchor-based label transfer functions: *FindTransferAnchors()* on SCT-normalized data with highly variable genes, followed by *TransferData()*.

To evaluate the concordance of the Reference cell annotations obtained with the COPILOT pipeline by Shahan et al. (2022), and that resulting from the SAPTICoN pipeline, we calculated the F1 score for each cell type. Cell identities were extracted from annotated Seurat objects, retaining only cells present in both data sets and excluding those with missing values. The predicted Reference and SAPTICoN labels were converted into factors with the same levels. A confusion matrix was generated with the confusionMatrix function of the caret package (Kuhn, 2008). Precision (positive predictive value) and recall (sensitivity) measures were extracted for each class and then used to calculate the F1 score per class according to the following formula: *F1=2×precision×recall/precision+recall*

We then calculated two global aggregations: (1) the macro F1-score, corresponding to the unweighted average of the F1-scores per class; (2) the weighted F1-score, calculated by weighting each score by the number of cells in the corresponding class (support). These measures indicate the overall accuracy of the SAPTICoN vs. Reference annotations (Figure 3F).

Metrics used to quantitatively assess the reciprocity of the Reference and SAPTICoN clustering data sets are the following: F1 score that balances precision (correct assignments) and recall (coverage of shared cells), providing an overall measure of agreement between data sets (Sasaki et al., 2008); Adjusted Rand Index (ARI) that quantifies the similarity between two clustering data sets, corrected for random chance (Hubert & Arabie, 1985); Entropy that measures the degree of disorder or uncertainty in cell assignments, with lower values indicating more consistent clustering (Shannon, 1948); Normalized Mutual Information (NMI) (Strehl & Ghosh, 2002) that assesses how much information is shared between the two clustering data sets, scaled between 0 (no overlap) and 1 (perfect match); Purity that evaluates the proportion of correctly matched cells within clusters, reflecting assignment consistency (Zhao & Karypis, 2001).

## SUPPLEMENTAL INFORMATION

Supplemental Figures S1 to S5, including legends. Seurat object.

## Author contributions

CP was involved in data production and curation, pipeline conceptualization, supervision, software development and analysis. He prepared the original draft and cowrote the manuscript. MV assisted with supervision, software development and infrastructure administration. AS initiated conceptualization, management and software development. GA assisted in software development. ED provided advise and critical assessment regarding methodology. PH initiated the project and its funding, participated to its conceptualization and management, and cowrote the manuscript.

## Acknowledgments

We wish to acknowledge Rachel Shahan, Che-Wei Hsu and coworkers who made available the Seurat object corresponding to their analysis of the Arabidopsis root cell atlas described in Shahan et al. (2022). The object matched the data presented in their study and was integrated as such in our work. We thank Joseph Tran for his generous support at the onset of the project and for in-depth discussions about algorithm design. We gratefully acknowledge the IPS2 Bioinformatics Platform for their assistance and support in providing computational resources and bioinformatics tools essential to this work.

---

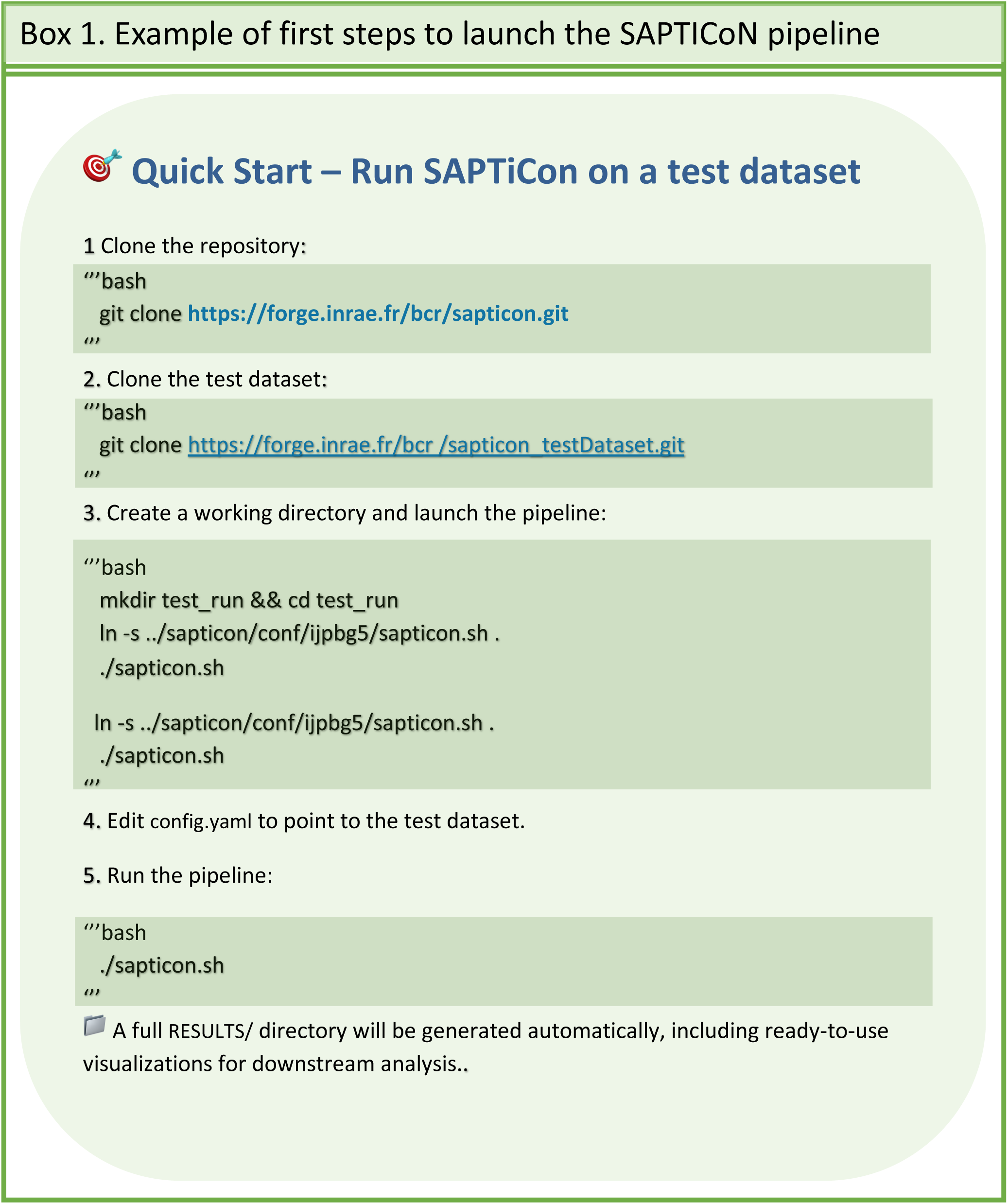

