## Supplemental Figures for "SAPTICoN, a robust no-code pipeline to analyze [plant] single cell transcriptomics data sets"

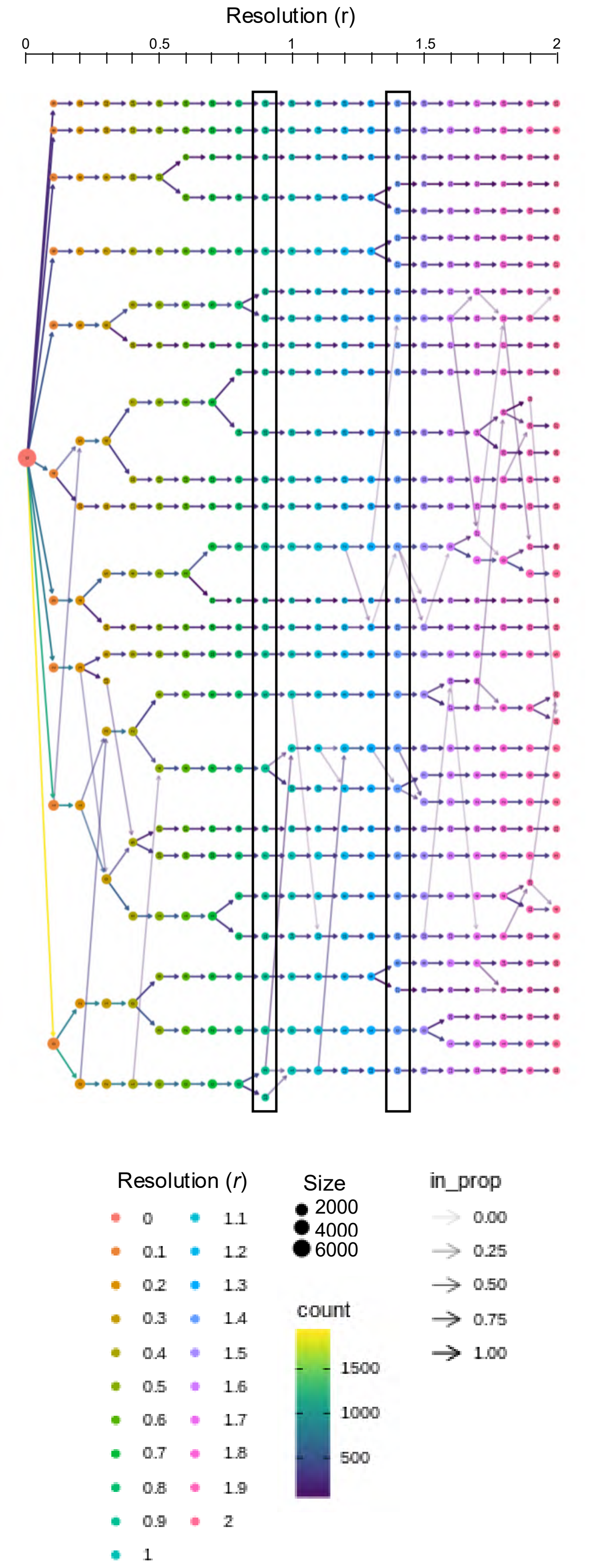


**Figure S1. Resolution tree for SAPTiCon clustering optimization.** The clustering tree of the analyzed data set of reference is shown with the dot size representing the number of cells associated to each cluster and the count parameter indicating the number of Seurat marker genes shared between cluster. Black boxes indicate the best resolution range to be considered.

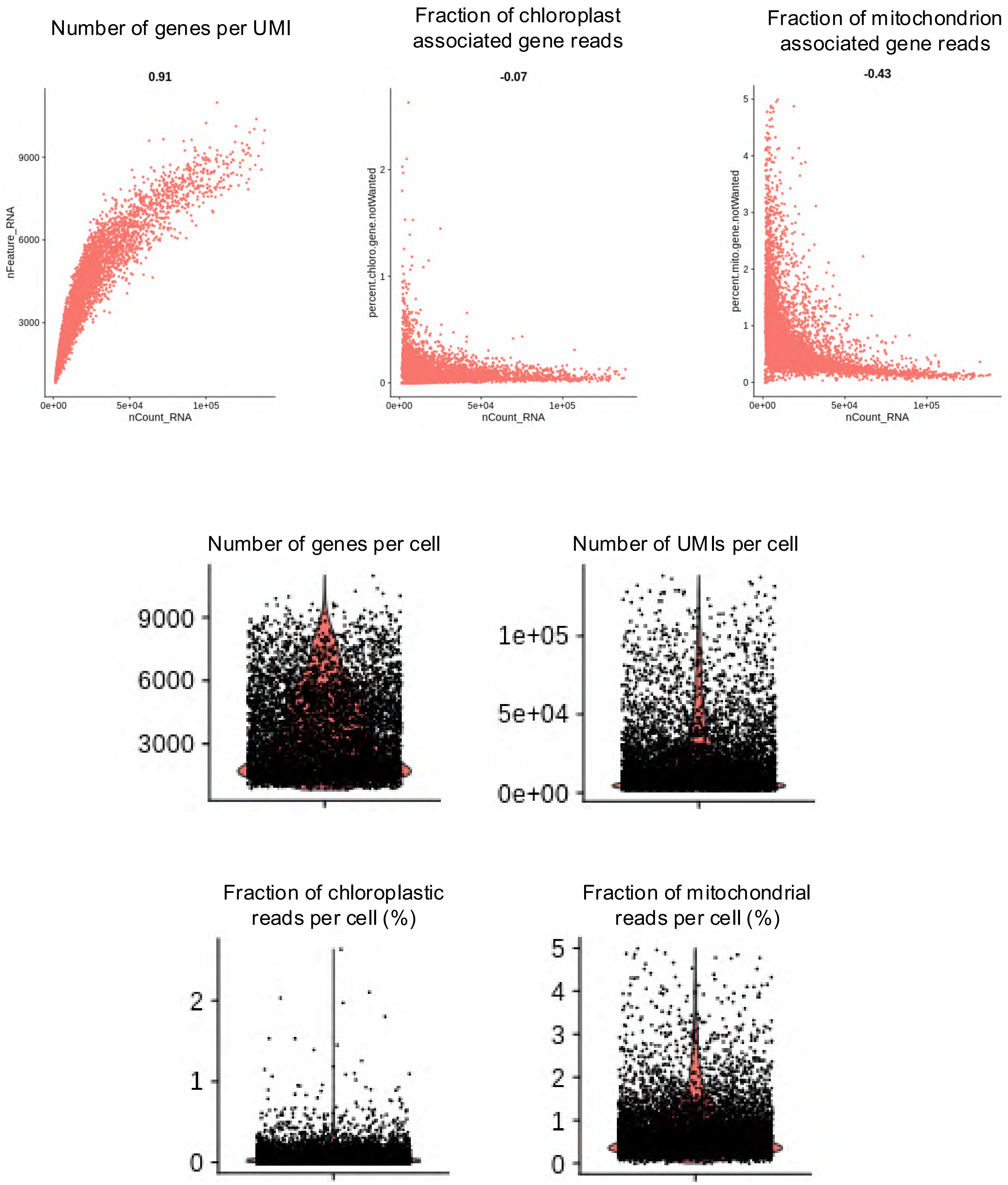


**Figure S2. Reference data quality control using SAPTiCoN implemented Seurat functions.**

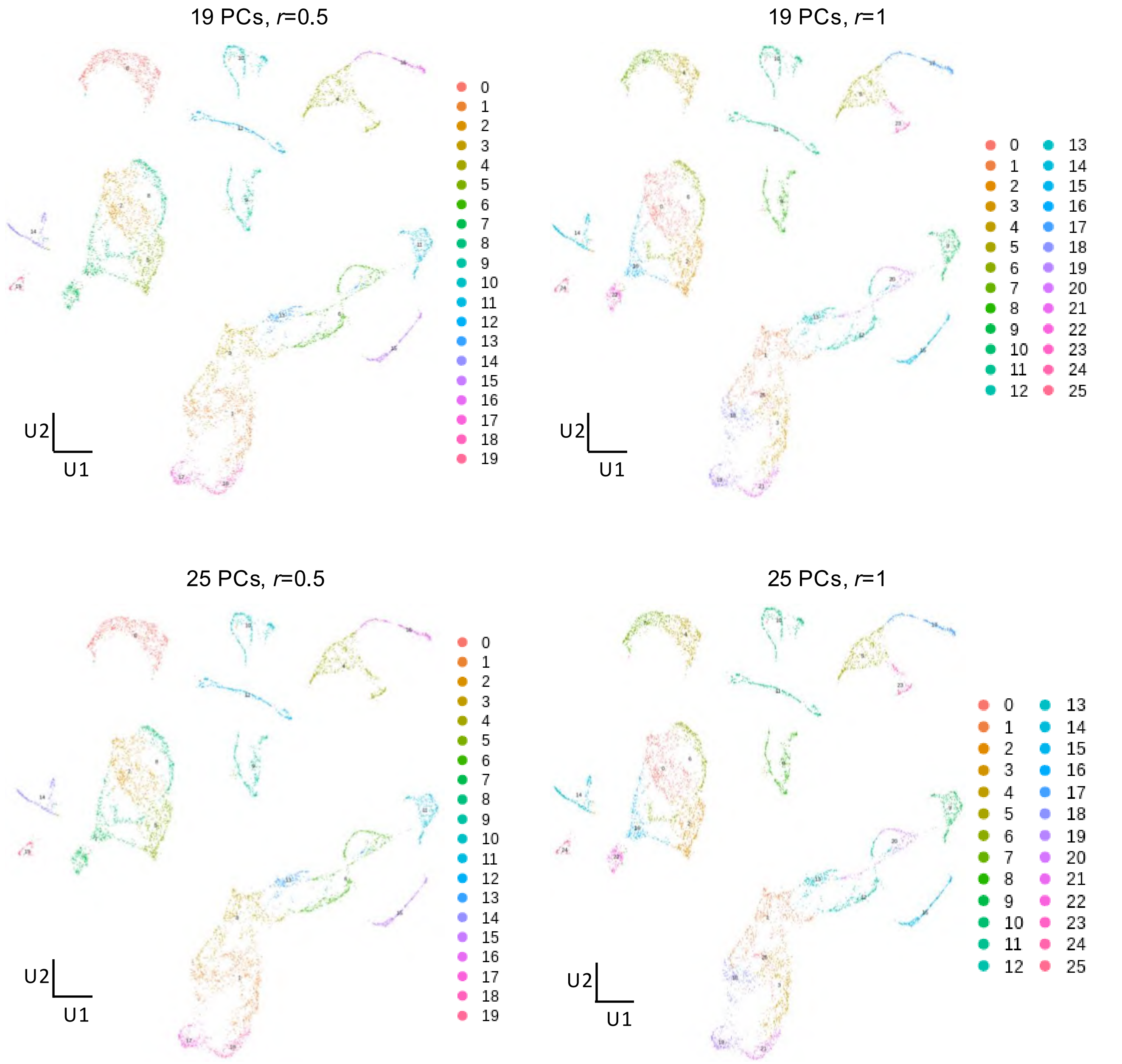


**Figure S3. Benchmark of SAPTICoN clustering according to the number of principal components (PC) and the resolution parameter (*r*).** UMAPs show the corresponding clustering results.
